# Frontal stimulation reshapes SCAN connectivity and improves motor function in Parkinson’s disease

**DOI:** 10.64898/2026.08.31.748359

**Authors:** Rupsha Panda, James A. Brissenden, Teresa Scerbak, Erin Proctor, Yiran Li, Michael Vesia, Roger L. Albin, Taraz G. Lee

## Abstract

Parkinson’s disease is a neurodegenerative disorder with characteristic motor deficits of bradykinesia and impaired coordination, and non-motor deficits affecting cognitive abilities such as attention and executive functions. Recent evidence suggests that some canonical motor deficits in Parkinson’s disease (e.g., freezing of gait) are associated with abnormalities in frontal cortical processing typically thought to underlie executive functions such as cognitive control. The evidence from human research linking frontal cognitive disruption and motor deficits in people with Parkinson’s disease is correlational. Individuals with cognitive impairments often also display gait and balance deficits, but causal evidence remains lacking. We combined non-invasive brain stimulation with functional MRI (fMRI) to ascertain whether dorsolateral prefrontal cortex (DLPFC) exerts a causal influence modulating motor function and to test the hypothesis that an excitatory stimulation protocol applied to a cognitive control network can improve motor performance in Parkinson disease.

Participants with Parkinson’s disease performed a precision force-tracking task shown to track with gait impairment while undergoing functional neuroimaging scans. On half of trials, participants were required to simultaneously perform a cognitively demanding 2-back working memory task to tax attentional systems. Following the initial baseline scan, participants returned for three separate counterbalanced sessions that combined transcranial magnetic stimulation (TMS) in the form of theta burst stimulation (TBS) with fMRI. Just prior to task performance, participants received either excitatory (intermittent TBS) or inhibitory (continuous TBS) stimulation protocols over right DLPFC. In the third session, participants received stimulation to a control site outside the cognitive and motor areas implicated in our task. Our within-subjects experimental design allowed us to examine the effects of stimulation on motor behavior, the influence of cognitive load on motor behavior, and the neural systems supporting cognitive control and motor execution.

Consistent with our hypothesis, excitatory stimulation of DLPFC improved motor performance in a precision force-tracking task as indexed by reduced tracking error and smoother movement execution. These effects emerged preferentially under dual-task conditions when cognitive demands were high. Our results show that this improved motor performance coincides with reductions in functional connectivity not only between the stimulation site and primary motor cortex, but also within the recently described somato-cognitive action network (SCAN) thought to support action planning and whole-body movement coordination.

We conclude that frontal cognitive control systems play a causal role in modulating movement in Parkinson’s disease and suggest that targeting these systems with non-invasive interventions may enhance motor function.

## Introduction

Parkinson’s disease (PD) is a neurodegenerative disorder historically characterized by progressive dopaminergic cell loss and accompanying motor dysfunctions. Although conceived as a disorder of the motor system, PD is a complex multisystem syndrome involving an array of motor and non-motor deficits.^1^ Cognitive impairments are virtually universal in PD, spanning a spectrum from subjective reports to overt dementia, usually coincident with disease progression.^2^ Mild to moderate PD is generally characterized by executive and attentional function deficits, even in individuals without cognitive symptoms.^3^ Convergent evidence indicates important interactions between cognitive and motor system deficits. Control of gait and ability to move seamlessly through the environment depends upon integration of sensory cues, motor commands, and cognitive demands, all filtered by attentional processes. Taxing working memory and attention with a concurrent secondary task, for example, impairs motor performance in both healthy individuals^4,5^ and those with Parkinson’s disease.^6–9^

Consistent with this cognitive-motor integration account, recent evidence strongly suggests that some canonical motor deficits in PD (e.g., freezing of gait) are associated with abnormalities in frontal cortical processing typically associated with executive functions such as cognitive control.^10–12^ These deficits may reflect disruptions of cognitive and motor networks linking basal ganglia, thalamic, and frontal cortical systems involved in the cognitive aspects of movement – an Attentional-Motor Interface.^13,14^ Recent work further suggests that a somato-cognitive action network (SCAN) involving strong connectivity between inter-effector regions of motor cortex and cognitive control networks supports action planning and whole-body movement coordination.^15^ Increases in connectivity within the SCAN may be a pathophysiologic signature of individuals with PD.^16^ These findings converge to suggest that frontal cognitive control systems are involved in supporting movement in PD, potentially compensating for progressive motor circuit dysfunction.

Despite the accumulation of evidence linking cognitive control dysfunction to motor impairment in Parkinson’s disease, the vast majority of existing evidence remains correlational. Individuals with cognitive impairments frequently exhibit gait and balance disturbances.^17–19^ Freezing of gait and other motor dysfunctions are associated with abnormal connectivity between cortical cognitive control networks and subcortical structures as measured by functional neuroimaging.^9,16,20–22^ Existing studies cannot establish whether alterations in activity of frontal cognitive control nodes play a causal role in supporting motor performance, reflect a compensatory response to motor dysfunction, or are simply a marker of an independent deficit that tracks with disease severity. Experimental manipulation of these networks provides an opportunity to determine whether frontal cognitive control systems causally contribute to motor behavior and represent viable therapeutic targets in Parkinson’s disease.

Transcranial magnetic stimulation (TMS) provides a means of directly testing the causal contribution of frontal cognitive control systems to motor behavior.^23,24^ TMS can transiently alter neuronal excitability within circumscribed cortical regions,^25–28^ enabling direct tests of the functional relevance of targeted brain networks. Repetitive stimulation protocols, such as theta burst stimulation (TBS), can induce relatively durable changes in cortical excitability following brief stimulation periods,^29^ enabling researchers to characterize behavior and brain activity in the time periods following stimulation. TMS protocols producing lasting effects have additionally shown promise as a therapeutic intervention across a range of neurological and psychiatric disorders, including PD.^30–32^

We combined functional neuroimaging with both excitatory and inhibitory TBS protocols to investigate the contribution of dorsolateral prefrontal cortex (DLPFC), a key node within cognitive control networks,^33,34^ to motor function in individuals with Parkinson’s disease. Immediately following stimulation, participants were asked to perform a precision force-tracking task shown to track with gait impairment^35,36^ while undergoing functional magnetic resonance imaging (fMRI). This design allowed us to examine the effects of stimulation on motor behavior, the influence of cognitive load on motor behavior, and explore the neural systems supporting cognitive control and motor execution. We hypothesized that stimulating DLPFC would not only alter the functional organization of cognitive-motor networks involved in movement control, but that excitatory stimulation would enhance motor performance.

## Materials and methods

### Participants

10 right-handed participants (6 females, 4 males; mean age: 65.2 years) were recruited locally from the University of Michigan and surrounding community. Participants met Movement Disorder Society (MDS) criteria for Clinically Probable Parkinson’s Disease.^37^ All participants were receiving stable dopaminergic medication at the time of testing and had taken their usual dose of medication prior to each session. Clinical and demographic characteristics are summarized in Table 1. Participants gave informed consent to participate. They were compensated $50 at the first study visit and were paid an additional $200 for completing the entire experiment. All participants underwent TMS Adult Safety Screening and fMRI Safety Screening to assess the potential risk of adverse reactions to TMS and MRI scanning. All experimental protocols were approved by the Institutional Review Boards of the University of Michigan Medical School (IRBMED #HUM00203427).

### Features of Parkinson’s disease participants at baseline

**Table 1.** Patient Demographics. . UPDRS = Unified Parkinson’s Disease Rating Scale; PD-CRS = Parkinson’s Disease – Cognitive Rating Scale; H&Y = Hoehn and Yahr Scale; GDS = Geriatric Depression Scale; LEDD = levodopa equivalent daily dose; SD = standard deviation;

| Feature | N | Mean | Range | SD |
| --- | --- | --- | --- | --- |
| Participants, N (Male / Female) | 10 (4 / 6) | – | – | – |
| Age | – | 65.2 | 56-72 | – |
| UPDRS Total | – | 36.6 | 22-68 | 15.3 |
| PD-CRS | – | 110.6 | 74–154 | 23.4 |
| H&Y | – | 1.9 | 1-3 | .70 |
| GDS | – | 5.7 | 0 – 13 | 4.4 |
| LEDD (mg/day) | – | 425.5 | 150–700 | 137.3 |

### Design Overview

We employed a within-subject design in which participants completed five separate sessions combining behavioral testing, fMRI, and TMS (Figure 1). In the initial session, all participants completed TMS and fMRI screenings^38^ and a baseline assessment. Participants were characterized using standard clinical and neuropsychological scales, including measures of parkinsonism (Movement Disorder Society–Unified Parkinson Disease Rating Scale [UPDRS]), mood (Geriatric Depression Scale [GDS]), apathy (Lille Apathy Rating Scale), and cognition using a battery of selected domain-specific tests.^22^ Bradykinesia was quantified using a tapping task that has been used extensively in prior studies.^39,40^

**Figure 1.**
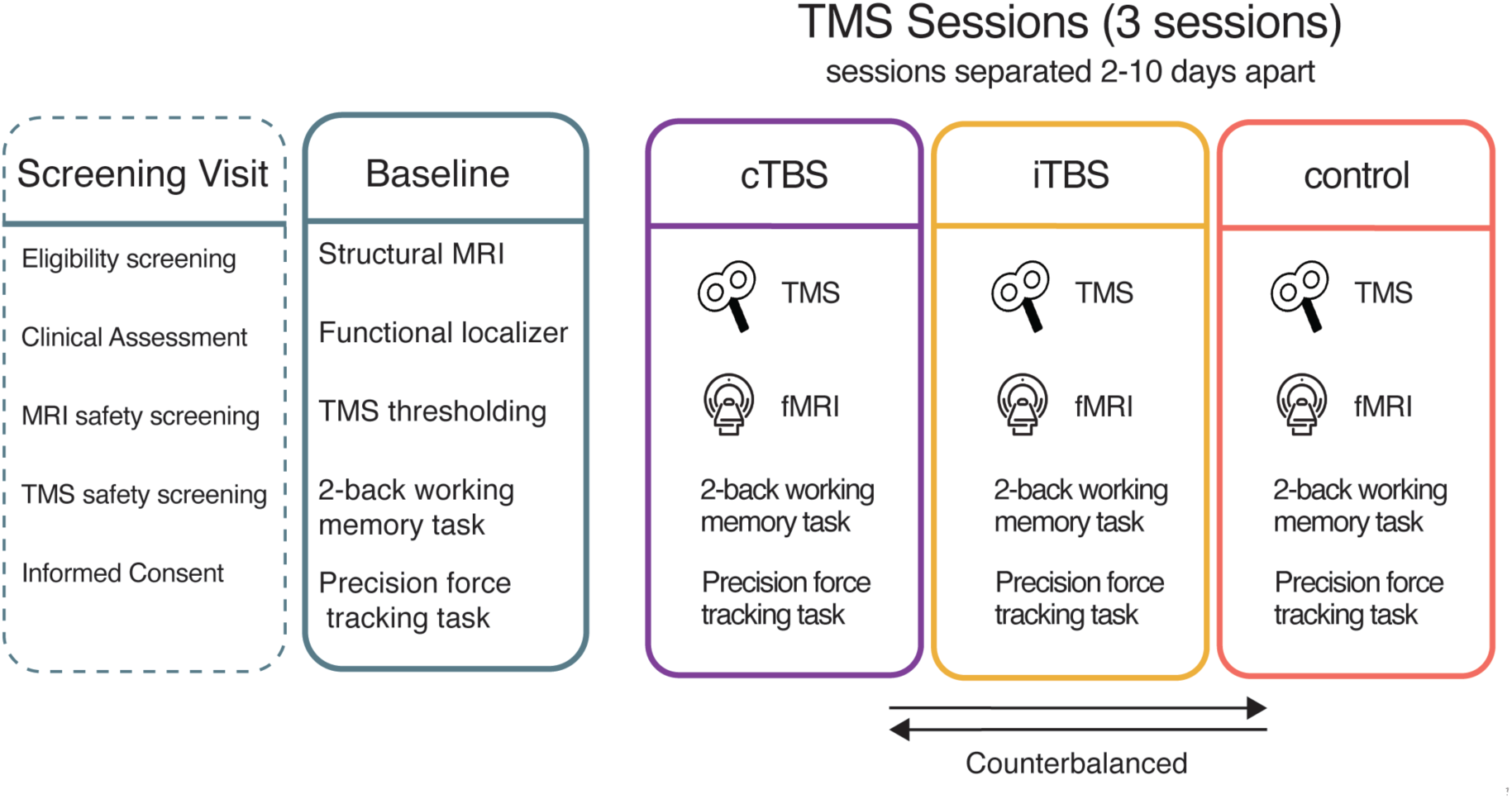
Experimental Timeline. Participants completed five study visits consisting of a screening visit, a baseline MRI session, and three combined TMS-fMRI experimental sessions. During the screening visit, participants underwent clinical, neuropsychological, and TMS safety assessments. During the baseline MRI session, structural and functional MRI data were acquired while participants performed the precision force-tracking task, and individualized DLPFC stimulation targets were identified from task-based fMRI. During each of the three subsequent experimental sessions, participants received either an excitatory stimulation protocol (intermittent theta burst stimulation; iTBS) over DLPFC, an inhibitory stimulation protocol (continuous theta burst stimulation; cTBS) over DLPFC, or stimulation over a control region (putative hip/leg area of somatosensory cortex) in a counterbalanced order. Immediately following stimulation, participants completed fMRI while performing the precision force-tracking task with and without concurrent 2-back working memory demands. Experimental sessions were separated by at least two days.

On a separate day, participants complete an MRI session to acquire structural scans, baseline measurements of resting-state functional connectivity, and task-based activation and functional connectivity. Frontal cortex stimulation targets were individually defined based on task-based measures derived from this MRI session (see below). A TMS assessment then determined each participant’s individual stimulation intensity that was used in the subsequent TMS-fMRI experimental sessions (see below). Participants returned for three separate combined TMS-fMRI sessions to assess the impact of prefrontal cortex perturbation on motor performance and brain network function. In these sessions, participants received either an excitatory stimulation protocol over DLPFC, an inhibitory stimulation protocol over DLPFC, or stimulation over a control site (see TMS Stimulation Sites). This control session allows for inferences to be made about the regional specificity of stimulation while also controlling for other non-specific effects of stimulation (auditory sounds, scalp sensation, etc.). Immediately following stimulation, participants walked across the hall and entered the fMRI scanning bay (average time to first functional scan: ∼8 minutes). In the scanner, participants completed a demanding precision-grip force task with and without cognitive load (see Precision force-tracking task). These sessions were spaced at least two days apart to prevent any carry-over effects, and the order of stimulation was counterbalanced across participants.

### Two-back working memory task

As a measure of executive function, participants performed a two-back working memory task at the beginning of each session. The task requires multiple cognitive abilities including attentional vigilance, short-term storage, and information manipulation.^41–43^ A centrally presented circle on a computer screen changed color once per second for 18 s, with a 500 ms inter-stimulus interval. Participants were instructed to respond at each color change whether the current color matched the color presented two trials earlier. Responses were made using a scanner-compatible force transducer (Current Designs, Inc., Philadelphia, PA), with one hand indicating a “yes” response and the other hand indicating a “no” response; response mapping was counterbalanced across participants. Accuracy rate was used as the dependent variable of interest. Each block comprised six trials, with a 6 s inter-block interval.

### Precision force-tracking task

To measure dynamic motor function in the scanner, participants performed a precision force-tracking task (FTT). Prior work has demonstrated that precision grip tasks track well with both disease progression and gait dysfunction in people with Parkinson’s disease.^35,36^ In this task (Figure 2), participants used a scanner-compatible force transducer to continuously modulate their grip force to match a target force output. Grip force was mapped onto a white cursor presented on the screen along the vertical dimension. Grip force was mapped onto the vertical position of a white cursor on the screen, such that zero force corresponded to the bottom of the display and thirty percent of each participant’s maximum squeeze force corresponded to the top of the display. Participants were instructed to keep the cursor within the bounds of a moving target circle. The target moved along the vertical axis according to a pre-generated trajectory formed from a linear combination of sinusoidal waveforms, producing a smooth, continuously varying force-tracking demand over the 11.7s tracking period. Target and cursor positions were updated and recorded on each screen refresh.

**Figure 2.**
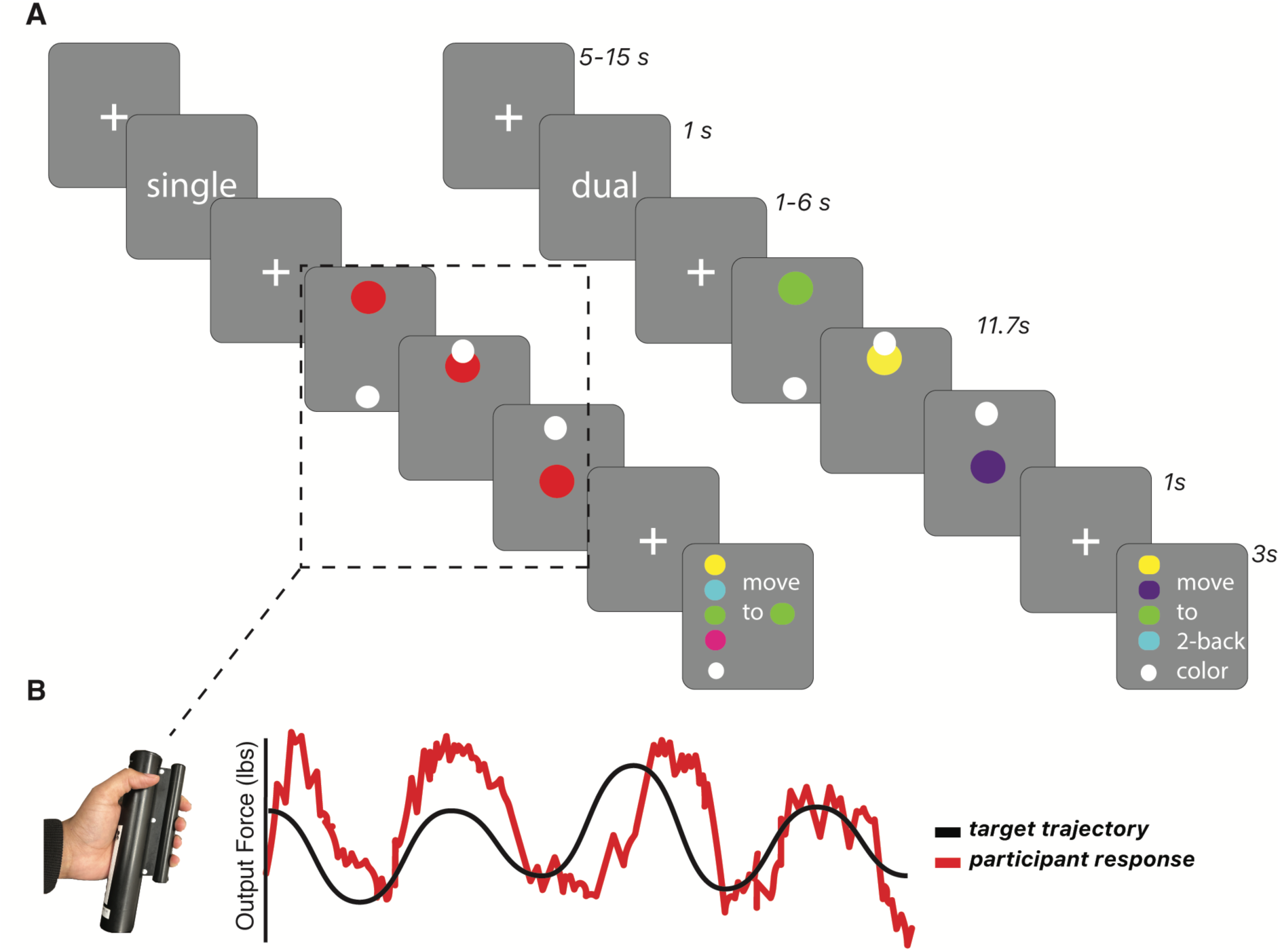
Precision force-tracking task performed during fMRI. **(A)** Trial structure for the single- and dual-task conditions. Each trial began with a task cue (“Single” or “Dual”), followed by a variable fixation interval and a 12s precision force-tracking period. During force tracking, participants continuously modulated grip force using a scanner-compatible force transducer to control the vertical position of a white cursor. Participants were asked to maintain this cursor within a target circle which moved vertically according to pseudorandom transformations of a sinusoidal waveform (see Methods). In dual-task trials, participants simultaneously performed a two-back working memory task in which the target circle changed color (∼0.5-0.75 Hz) throughout the tracking period. At the end of the tracking period, participants reported the color presented two color changes prior to the final color by using the force transducer to select one of the four colors presented on the screen. To match these added visuomotor demands on single-task trials, participants were cued to select one of four colors presented at the end of each single-task tracking period. **(B)** Example force-tracking performance on a single trial. The black trace represents the target force trajectory and the red trace represents participant-generated grip force.

To manipulate cognitive demands during motor performance, half of the trials involved force tracking alone (single-task condition), whereas the remaining half required participants to simultaneously perform a working memory task during force tracking (dual-task condition). Single- and dual-task trials were presented in a randomized order, with each block containing four trials of each condition. Each task block consisted of eight trials, with a total of 80 trials completed across the experiment.

During dual-task trials, participants performed the force-tracking task while simultaneously completing a 2-back task. The target circle changed color pseudo-randomly 5–9 times per trial, requiring participants to track the target while monitoring the color sequence. At the end of each trial, participants used the grip-force sensor to identify the color presented two color changes before the final color displayed (Fig. 2A “dual”). The pace of changing colors differed from trial to trial, such that participants had to hold every 2-back color within their working memory throughout the trial. This manipulation required both tonic attentional vigilance and working memory maintenance, two executive functions previously implicated in attentional-motor integration.^44,45^ Taxing executive function using dual-task paradigms has been shown to interfere with motor function in people with Parkinson’s disease.^6,8,17^ To match low-level visuomotor demands between conditions, at the end of each single-task trial four colored circles were presented in the same configuration as at the end of dual-task trials and participants were instructed to move the cursor to a specified target circle. This ensures that the fMRI contrast between single- and dual-tasking conditions reflect dual-task cognitive demands rather than any activity associated with the 2-back response period.

### Image acquisition and processing

#### MRI acquisition

MR data were acquired with a GE Signa MR750 3.0T MRI scanner with a 32-channel head coil. The baseline session collected high-resolution 3D inversion-recovery T1-weighted MPRAGE (sagittal; TR = 2523 ms; TE = 3.55 ms; TI = 1060 ms; flip angle = 8°; 0.8 mm isotropic voxels; matrix = 320 × 320; 208 slices; parallel imaging: auto-calibrating reconstruction for cartesian imaging (ARC) 2x in-plane and 2x through-plane) and 3D CUBE T2-weighted scans (sagittal; TR = 5202 ms; TE = 60 ms; flip angle = 90°; 0.8 mm isotropic voxels; matrix = 320 × 320, 226 slice; parallel imaging: ARC 2x in-plane and 2x through-plane). Functional data for resting-state and task scans were acquired with a 2D gradient-echo echo-planar imaging (EPI) sequence using simultaneous multi-slice acceleration (GE HyperBand; MB = 8): TR = 650 ms; TE = 35 ms; flip angle = 52°; 2.3 mm isotropic voxels; matrix = 92 x 92; 64 slices with 0% inter-slice gap. Spin-echo EPI field maps were also acquired with opposite phase encoding directions (Anterior-to-Posterior; Posterior-to-Anterior) and geometry matched to the functional EPI timeseries. Ten functional runs of 5 min each were collected.

#### MRI preprocessing

Anatomical and functional data were preprocessed using *fMRIPrep*^46^ (Version 20.2.5; RRID:SCR_016216). The T1-weighted (T1w) image was corrected for intensity non-uniformity with *N4BiasFieldCorrection*,^47^ distributed with ANTs 2.3.3^48^ (RRID:SCR_004757), and used as the T1w-reference throughout the workflow. The T1w-reference was then skull-stripped with a *Nipype* implementation of the *antsBrainExtraction.sh* workflow (from ANTs), using OASIS30ANTs as target template. Brain tissue segmentation of cerebrospinal fluid (CSF), white-matter (WM) and gray-matter (GM) was performed on the brain-extracted T1w using *fast*^49^(FSL 5.0.9; RRID:SCR_002823). Brain surfaces were reconstructed using *recon-all*^50^ (FreeSurfer 6.0.1; RRID:SCR_001847), and the brain mask estimated previously was refined with a custom variation of the method to reconcile ANTs-derived and FreeSurfer-derived segmentations of the cortical gray-matter of Mindboggle^51^ (RRID:SCR_002438). Freesurfer outputs were then input into *ciftify_recon_all*, which performs surface-based alignment using the MSMSulc algorithm^52^ and resamples data from native space to 32k fsLR space. The fsLR 32k space combines the benefits of nonlinear MNI volumetric registration (for subcortical structures; FSL FNIRT) with surface-level registration based on sulcal anatomy for the cortex.^53^

For each BOLD run, the following preprocessing was performed. First, a reference volume and its skull-stripped version were generated using a custom methodology of *fMRIPrep*. A B0-nonuniformity map (or *fieldmap*) was estimated based on two (or more) echo-planar imaging (EPI) references with opposing phase-encoding directions, with *3dQwarp*^54^ (AFNI 20160207). Based on the estimated susceptibility distortion, a corrected EPI (echo-planar imaging) reference was calculated for a more accurate co-registration with the anatomical reference. The BOLD reference was then co-registered to the T1w reference using *bbregister* (FreeSurfer).^55^ Co-registration was configured with six degrees of freedom. Head-motion parameters with respect to the BOLD reference (transformation matrices, and six corresponding rotation and translation parameters) were estimated before any spatiotemporal filtering using *mcflirt*^56^ (FSL 5.0.9). The BOLD time-series were resampled onto their original, native space by applying a single, composite transform to correct for head-motion and susceptibility distortions. Gridded (volumetric) resamplings were performed using *antsApplyTransforms* (ANTs), configured with Lanczos interpolation to minimize the smoothing effects of other kernels.^57^ For the definition of TMS target regions, resting-state timeseries were analyzed in native T1 space. These native T1w timeseries were smoothed in a parcel-constrained manner (cortical ribbon and subcortical gray matter structures) with a 4mm FWHM kernel to maximize spatial accuracy of the TMS target definitions. For task-based functional connectivity analysis, this smoothing was done after additional preprocessing was performed (see Task-based functional connectivity preprocessing below).

#### Task-Based Functional MRI processing

Following fMRIPrep, we performed confound regression on the task-based timeseries using the *nilearn* python package (Version 0.8.1; RRID:SCR_001362; https://doi.org/10.5281/zenodo.8397156). Several confounding time-series were calculated based on the preprocessed BOLD: framewise displacement (FD) and three region-wise global signals. FD was computed using the Power formulation^58^. The three global signals were extracted within the CSF, the white matter, and the whole-brain masks. The confound time series derived from head motion estimates and global signals were expanded with the inclusion of temporal derivatives and quadratic terms for each.^59^ Frames that exceeded a threshold of 0.2 mm FD were considered motion outliers. The confound regression included the six head motion estimates and three region-wise global signals (CSF, WM, whole-brain), their temporal derivatives, quadratic terms, and the quadratic expansions of their derivatives (36 regressors).^59^ The first 16 volumes of each functional run were removed prior to modeling to allow for signal equilibrium. The resulting residual timeseries were *z*-scored. Functional connectivity analyses were performed on residual time series obtained after regressing out task-evoked activity using a finite impulse response (FIR) GLM in *nilearn* (version 0.11.1). The FIR GLM was performed at each cortical vertex and subcortical voxel, after which residual time series were extracted from predefined regions of interest (ROI). For each participant, ROI-to-ROI functional connectivity was quantified using Pearson correlations between residual time series, and correlation coefficients were Fisher z-transformed prior to group-level statistical analyses.

#### Regions of Interest

Regions of interests (ROI) were defined using multiple atlases. Cortical ROIs were derived from the Human Connectome Project multimodal parcellation (Glasser atlas). The dorsolateral prefrontal cortex ROI consisted of areas p9-46v, 46, 9-46d, and a9-46v, while motor cortical ROIs included area 4. Striatal ROIs were derived from the Oxford-GSK-Imanova Striatal Connectivity Atlas using the sensorimotor (putamen) and executive (caudate) subdivisions in this parcellation. Somato-cognitive association network (SCAN) ROIs were derived from the Human Connectome Project motor task data. Hand, foot, and mouth motor representations were identified using the MOTOR contrasts (LF-AVG, LH-AVG, RF-AVG, RH-AVG, and T-AVG). These maps were thresholded at z > 2. Thresholded maps were then constrained to the primary motor strip from the Glasser atlas. Hand, foot, and mouth representations from the HCP data were masked out to isolate inter-effector sensorimotor regions. These resulting ROIs were used in subsequent ROI-to-ROI functional connectivity analyses examining connectivity between SCAN regions, DLPFC, motor cortex, and striatal subdivisions.

### Transcranial Magnetic Stimulation

#### Protocol

TMS was delivered using a MagPro X100 magnetic stimulator with a 70-mm figure-8 coil (MC-B70, MagVenture Inc.). To determine individualized stimulation intensity, we assessed motor excitability following the baseline fMRI session. Motor evoked potentials (MEP) were elicited using biphasic pulses and were recorded from the right first dorsal interosseous (FDI) muscle using surface electromyography (2-channel EMG device, Rogue Research, Montreal). Active motor threshold (AMT) was defined as the minimum percentage of stimulator output that elicited an MEP of at least 50 μV peak-to-peak amplitude on five of ten trials while the participant maintained a 20% maximum voluntary contraction. At the end of the thresholding session, participants were given several pulses at 80% AMT over regions of the scalp roughly corresponding to the DLPFC and the control site to ensure that the stimulation intensity would be tolerable. In the TMS-fMRI sessions, we delivered theta-burst stimulation (TBS) at 80% AMT using standard parameters, consisting of bursts of three pulses at 50 Hz repeated every 200 ms. Continuous TBS (cTBS) consisted of 600 pulses over 40 seconds. Intermittent TBS (iTBS) was applied in two second trains repeated every ten seconds for a total of 600 pulses over 190 seconds.^29^ cTBS decreases motor cortical excitability, whereas iTBS increases cortical excitability for up to approximately 60 min following stimulation.^29^ We have shown previously that TBS protocols can be used to modulate prefrontal cortical function and large-scale brain networks.^26,60–62^

#### Stimulation Site

Stimulation targets were individually defined based on analyses of anatomical and task-based data from the baseline fMRI session. To identify individualized DLPFC stimulation targets, task-based fMRI data acquired during the baseline session were analyzed using a GLM implemented in AFNI. Neural responses were modeled using four task regressors: single-task cue, dual-task cue, single-task tracking, and dual-task tracking periods. Event onsets were convolved with AFNI’s duration-modulated block response function (dmBLOCK). The GLM was estimated using 3dDeconvolve followed by 3dREMLfit to account for temporal autocorrelation in the fMRI time series. Subject-specific DLPFC targets were defined using a dual-task versus single-task force tracking contrast (dual > single). This contrast identified brain regions exhibiting greater activation during the dual-task condition, which combined cognitive load with precision force-tracking, relative to motor task performance alone (Figure 3). An individualized DLPFC stimulation target was defined for each participant as the peak activation within this contrast map constrained to an anatomically defined mask of the right middle frontal gyrus. A TMS neuronavigational system (Brainsight 2, Rogue Research) aligned the previously acquired structural scan to each participant’s skull in real time to allow for precise and consistent targeting across all sessions. To control for nonspecific effects of TMS, including sensory stimulation induced current magnitude, an active control stimulation site was used for comparison with DLPFC stimulation. The control site was in the putative hip/trunk representation of primary somatosensory cortex (S1). This region was identified anatomically by first identifying right postcentral gyrus and then selecting a stimulation target five mm lateral from the midline. This region did not exhibit task-related activation in any participants in the present study. cTBS was applied to this region in all control sessions.

**Figure 3.**
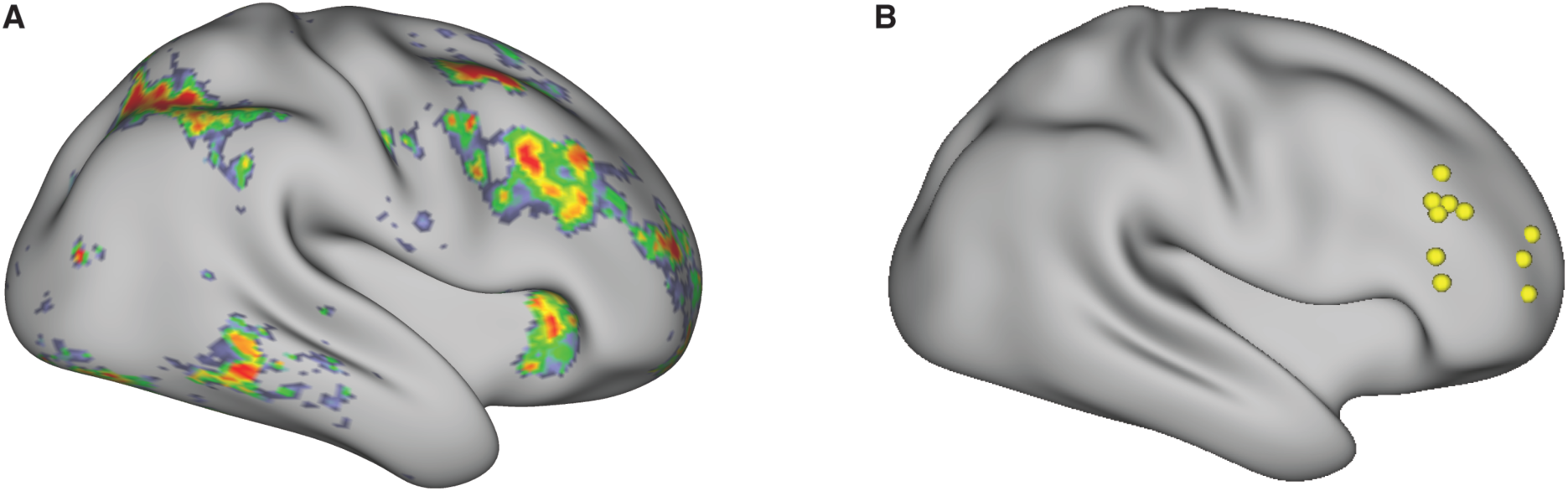
Individualized TMS target in dorsolateral prefrontal cortex. **(A)** Example contrast map of activation between dual-task and single-task blocks from the baseline session (thresholded at Z > 3 for visualization purposes). For each participant, the peak of the cluster of activation within middle frontal gyrus was selected as the TMS target. **(B)** Individualized TMS stimulation targets for all participants (yellow spheres) projected onto an inflated cortical surface. Neuronavigation software (Brainsight) was used to guide the position of the TMS coil to align with this target in each subsequent experimental sessions.

### General behavioral analysis

Behavioral data were analyzed using Bayesian hierarchical models implemented in the *brms* package in R. For each outcome, models included fixed effects of stimulation condition (TMSSite: iTBS, cTBS, control), task condition (dual vs. single, where applicable), and experimental day (to regress out order effects), along with a random intercept for subject. We calculated dependent measures of percent accuracy, root mean squared error (RMSE), and jerk in the force tracking task. Percent accuracy was defined as the percentage of frames (sampling rate = 60 Hz) during which the cursor remained within the target circle on each trial. Trialwise RMSE was calculated as the square root of the mean squared distance between the cursor position and the target position across all frames within a trial. The smoothness of motor performance was quantified using measure of jerk, which was estimated using successive discrete differences corresponding to the third derivative of force with respect to time after the force signal was interpolated to ensure uniform sampling.^63^ Mean squared jerk was then computed across each trial and normalized by movement duration, with lower values indicating smoother force production and higher values reflecting more abrupt fluctuations in force output. We did not evaluate snap, crackle, pop or other rate of change metrics in this data set. Models were estimated using four Markov chain Monte Carlo (MCMC) chains with 6000 iterations each, with the first 2000 iterations discarded as warm-up. Convergence was assessed using the potential scale reduction factor (R-hat)^64^ and effective sample size. Posterior predictive checks confirmed that model assumptions were satisfied. To facilitate interpretation, we additionally computed posterior distributions of pairwise differences between stimulation conditions within each task condition. Statistical analyses were performed within a Bayesian framework. Effects are summarized using the posterior median and 95% highest density interval (HDI), which represents the range containing the most credible values of the parameter estimate. A main advantage of Bayesian analyses is that it allows inferences to be made by evaluating the span of the entire posterior distribution. Nevertheless, we also report the probability of direction (pd) to aid in inference. This measure is the proportion of the posterior distribution that lies in the same direction as the posterior median. The pd ranges from 50% (no evidence for an effect) to 100% (all posterior samples support the same directional effect). We refer to pd ≥ 97.5% as strong evidence for an effect, 90% ≤ pd < 97.5% as moderate evidence, and pd < 90% as little evidence for an effect. We adopt theses thresholds for ease of communication in the text, but we must note that the choice of a hard threshold for inference is inherently arbitrary.

## Results

### Excitatory stimulation of DLPFC improves motor performance under increased cognitive demand

We first sought to assess whether non-invasive perturbation of DLPFC affects motor function during a precision force-tracking task. Motor performance was quantified using several different metrics: trial wise root mean squared error (RMSE), accuracy rate, and jerk (movement smoothness). RMSE quantifies the amount of deviation from optimal tracking performance over the course of a trial, where lower values reflect more accurate force tracking (Figure 4A). Motor performance following iTBS resulted in lower RMSE relative to performance following cTBS during both Single-task (median [HDI] = -0.006 [-0.011, -0.001], *pd* = 98.5%) and Dual-task conditions (median [HDI] = -0.015 [-0.022, -0.008], *pd* = 100%). iTBS also resulted in reduced RMSE during Dual-task performance relative to control site stimulation (median [HDI] = -0.014 [-0.021, -0.008], *pd* = 100%). We found moderate evidence that iTBS led to reduced RMSE during Single-task performance relative to control site stimulation (median [HDI] = -0.004 [-0.009, 0.001], *pd* = 92.7%) There was little evidence that cTBS differed from control stimulation in either condition (Single-task median [HDI] = -0.002 [-0.007, 0.003], *pd* = 77.1%; Dual-task median [HDI] = -0.001 [-0.007, 0.006], *pd* = 57.8%).

**Figure 4.**
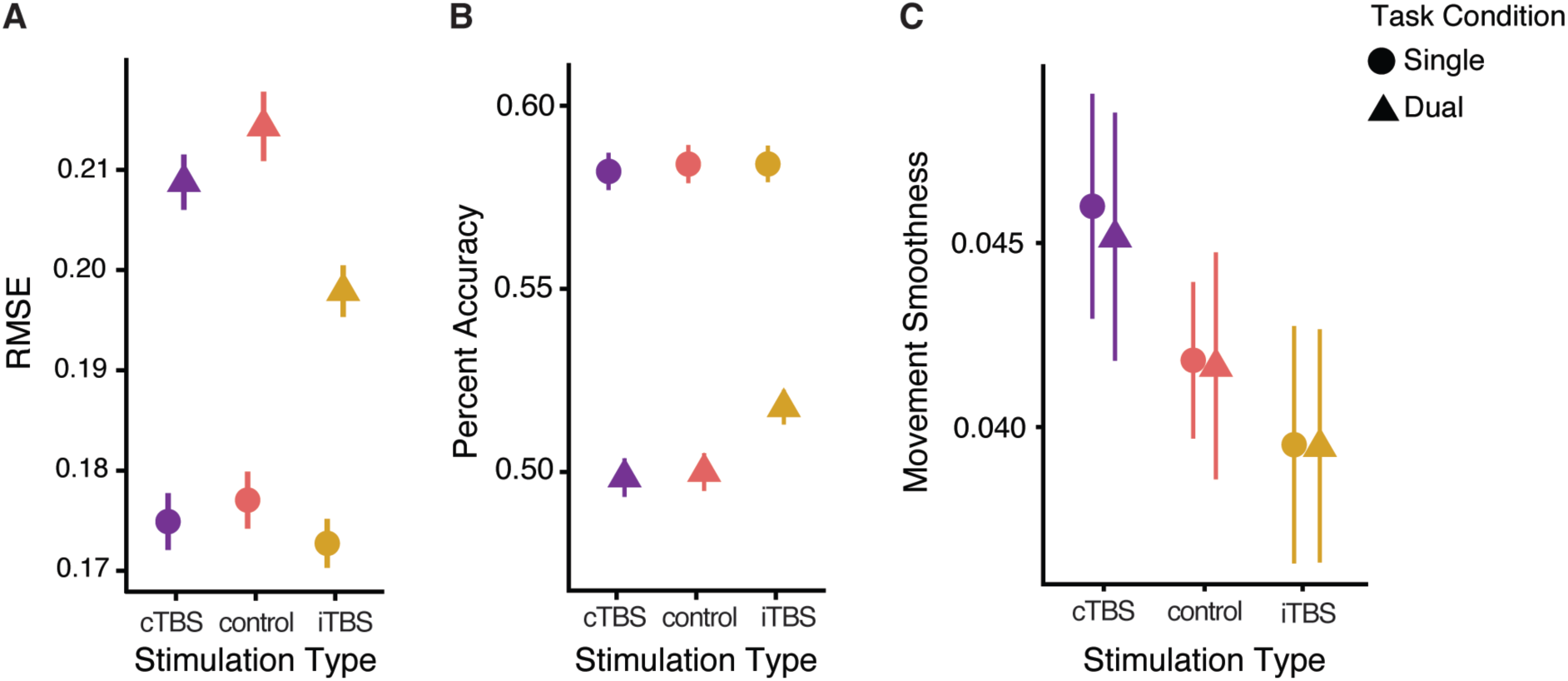
Excitatory stimulation over DLPFC improves motor performance. **(A)** Root mean squared error (RMSE) during precision force tracking across stimulation conditions (cTBS, control, and iTBS) and task condition (dual or single). Lower values of RMSE indicate more accurate tracking performance. **(B)** Percent accuracy, defined as the percentage of time the cursor remained within the target boundary. Higher values indicate better tracking performance. **(C)** Normalized jerk, a measure of movement smoothness, shown across stimulation and task conditions. Lower jerk values indicate smoother movement trajectories. Points represent group means and error bars denote ±1 SEM.

These results showing the benefit of excitatory DLPFC stimulation on motor performance were mirrored when examining the related dependent measure of accuracy rate. Accuracy rate was quantified as the percentage of frames in which the cursor remained within the target boundary, with higher values reflecting better tracking performance (Figure 4B). iTBS resulted in a higher accuracy rate relative to cTBS during Dual-task performance (median [HDI] = -2.29 [-3.66, -1.00], *pd* = 99.96%), with little evidence for differences in the Single-task condition (median [HDI] = -0.69 [-2.12, 0.88], *pd* = 81.4%). Similarly, iTBS also increased accuracy rate relative to control site stimulation during Dual-task performance (median [HDI] = -1.75 [-3.09, -0.46], *pd* = 99.7%). In contrast, there was little evidence for meaningful differences between cTBS and control stimulation (median [HDI] = 0.545 [-0.734, 1.87], *pd* = 79.4%).

Aggregate motor performance was reliably improved following iTBS relative to both cTBS and control site stimulation, but what kinematic features of action give rise to this improvement? Jerk is a measure of movement smoothness, where lower values reflect smoother and less abrupt force adjustments (Figure 4C). There was no evidence for differences in jerk between Dual and Single conditions (median [HDI] = -0.001 [-0.006, 0.003], *pd* = 68.1%), therefore, jerk analyses examining the effects of stimulation collapsed across these conditions. iTBS reduced jerk relative to cTBS (median [HDI] = 0.006 [0.0001, 0.012], *pd* = 98%), indicating smoother motor output following excitatory stimulation of DLPFC. There was little evidence for meaningful differences between iTBS and control stimulation (median [HDI] = .002 [-.004, .008], *pd* = 77.5%). There was also little evidence for meaningful differences between cTBS and control condition (median [HDI] = -0.0038 [-0.009, 0.002], *pd* = 90.2%).

iTBS produced reliable improvements in motor performance relative to both cTBS and control site stimulation, with effects emerging most strongly under dual-task conditions that taxed cognitive function. These improvements were evident across measures of RMSE, accuracy rate, and movement smoothness, suggesting that excitatory stimulation of DLPFC enhanced multiple aspects of motor control.

### Improved motor performance after iTBS cannot be explained by a tradeoff with cognitive performance

The most pronounced improvement of motor performance following excitatory stimulation to DLPFC was during dual-task conditions where participants were asked to simultaneously perform a cognitively demanding 2-back working memory task during force-tracking. One possibility is that motor performance improved not because of overall enhanced function, but because participants shifted priority to focus on motor performance at the expense of cognitive performance. We did not find this to be the case as cognitive performance was largely unaffected by stimulation, both when the 2-back task was performed alone and when it was combined with motor tracking. Bayesian analyses provided little evidence that stimulation altered cognitive performance in either context. During dual-tasking, there was weak evidence for *improved* 2-back accuracy following iTBS relative to cTBS (median difference = 0.050, 95% HDI [−0.021, 0.121], *pd* = 0.925) and *reduced* accuracy following cTBS relative to control site stimulation relative (median difference = 0.046, 95% HDI [−0.026, 0.119], *pd* = 0.905). 2-back performance was similar between iTBS and control stimulation (median difference = 0.004, 95% HDI [−0.069, 0.078], *pd* = 0.541). Performance on the standalone 2-back task completed prior to performance of the force-tracking task did not differ across stimulation conditions, with all pairwise comparisons exhibiting broad HDIs spanning zero and limited evidence for stimulation-related effects. These findings indicate that DLPFC stimulation produced relatively minimal changes in working memory performance. Motor benefits observed following iTBS were not due to a shift in priority away from cognitive task performance but rather reflect an enhanced capability to perform a complex motor task in the face of distraction.

### Excitatory stimulation of DLPFC ameliorates aberrant hyper-connectivity with primary motor cortex

Prior work has demonstrated that persons with Parkinson’s disease have elevated levels of functional connectivity between frontoparietal brain areas important for cognitive control and both cortical and sub-cortical motor regions.^16,20–22^ Behaviorally, excitatory stimulation delivered to DLPFC leads to improvements in motor performance, but how does this impact cognitive-motor network activity that supports performance?

We examined the effect of DLPFC stimulation on functional connectivity between primary motor cortex and DLPFC, which has been previously implicated in cognitive-motor integration (Figure 5). Functional connectivity was quantified as Fisher z-transformed Pearson correlations between the residual time series of each region after regressing out task-evoked activity. iTBS reduced DLPFC–M1 functional connectivity relative to cTBS (median [HDI] = -0.143 [-0.265, -0.016], *pd* = 98.7%), with weaker evidence for differences between iTBS and control stimulation (median [HDI] = -0.060 [-0.185, 0.064], *pd* = 84%). Likewise, there was little evidence that functional connectivity differed between cTBS and control site stimulation (median [HDI] = -0.083 [-0.203, 0.047], *pd* = 91.1%). Excitatory stimulation of DLPFC altered cognitive-motor network interactions, potentially reflecting reduced coupling within networks that have been shown to display pathological hyperconnectivity in Parkinson’s disease.

**Figure 5.**
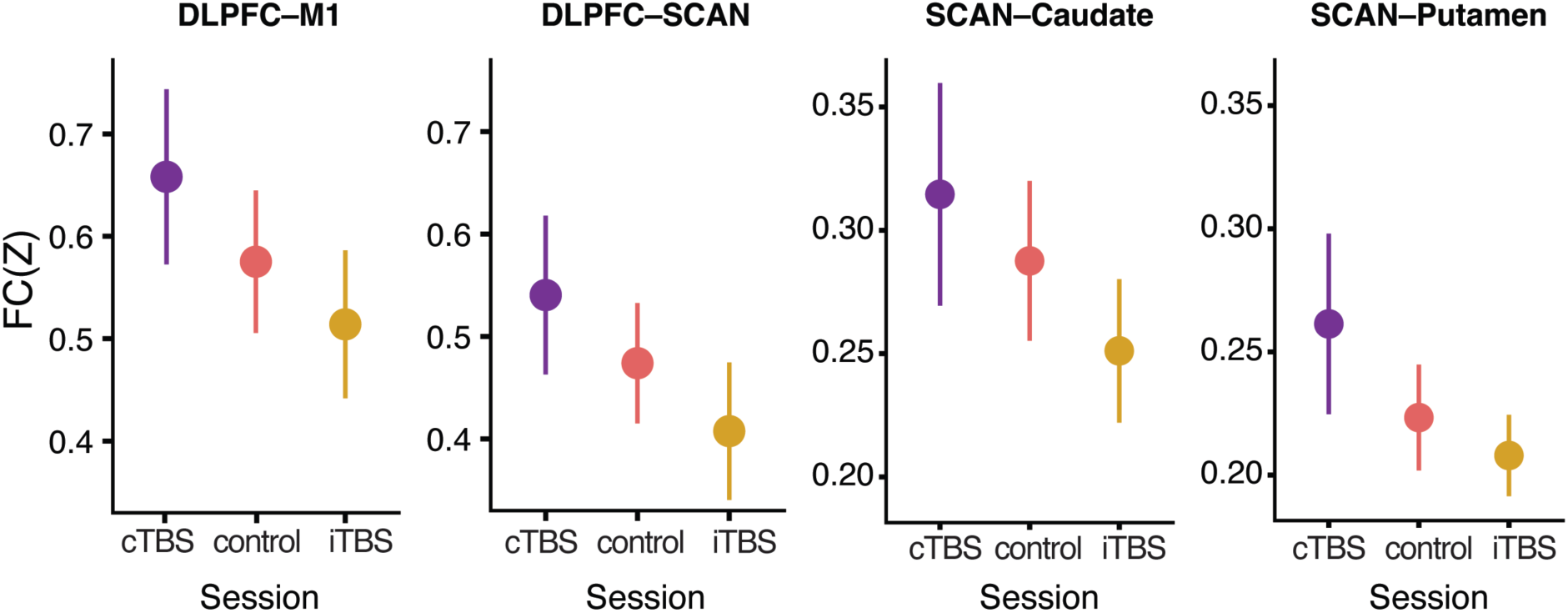
Excitatory stimulation of DLPFC alters functional connectivity within cognitive–motor networks. Functional connectivity was quantified as Fisher *z*-transformed Pearson’s correlations between regions using the residual BOLD time series from the task-based GLM. DLPFC regions of interest were defined individually from activation maps from each individual’s baseline scanning session intersected with an anatomical mask. Non-effector regions of motor cortex (somato-cognitive action network nodes; SCAN) and striatal regions of interest were defined from a combination of task-based and anatomical atlases (see Methos). From left to right, panels show connectivity between the following regions of interest across stimulation conditions: DLPFC–M1, DLPFC–SCAN, SCAN–caudate, and SCAN–putamen. Points represent group means and error bars denote ±1 SEM.

### Excitatory stimulation alters connectivity of somato-cognitive action network nodes in motor cortex

The previous analysis focused on primary motor cortex as a single unit. However, recent evidence suggests that motor cortex is composed of alternating effector-specific regions and inter-effector regions that integrate cognition and action through their connectivity with canonical attention networks.^15^ As this somato-cognitive action network (SCAN) has been shown to display hyperconnectivity with subcortical nuclei in Parkinson’s disease,^16^ we hypothesized that stimulation-induced benefits of DLPFC stimulation might coincide with changes to SCAN connectivity.

Stimulation-dependent differences emerged across several SCAN-centered networks. Relative to cTBS, iTBS reduced functional connectivity between SCAN nodes in motor cortex and DLPFC (median [HDI] = 0.133 [0.008, 0.249], *pd* = 98.4%), SCAN and caudate (median [HDI] = 0.064 [0.006, 0.123], *pd* = 98.2%), and SCAN and putamen (median [HDI] = 0.054 [-0.005, 0.111], *pd* = 96.6%). In contrast, comparisons involving control site stimulation were consistently smaller and more uncertain across all three networks (SCAN– DLPFC: control vs cTBS, median [HDI] = -0.067 [-0.189, 0.054], *pd* = 87.4%; control vs iTBS, median [HDI] = 0.066 [-0.054, 0.186], *pd* = 86.7%; SCAN–caudate: control vs cTBS, median [HDI] = -0.027 [-0.088, 0.030], *pd* = 82.3%; control vs iTBS, median [HDI] = 0.037 [-0.022, 0.094], *pd* = 90.2%; SCAN–putamen: control vs cTBS, median [HDI] = -0.038 [-0.099, 0.017], *pd* = 90.6%; control vs iTBS, median [HDI] = 0.015 [-0.044, 0.072], *pd* = 70.9%). Together, these findings demonstrate that the motor improvements observed following iTBS were accompanied by reduced connectivity across multiple SCAN-centered cognitive-motor networks.

Across all functional connectivity analyses, the behavioral improvements observed following iTBS were accompanied by coordinated changes in functional connectivity among DLPFC, SCAN, and striatal regions. Thus, these effects were evident across multiple cortical and cortico-striatal networks implicated in the cognitive control of movement

## Discussion

Accumulating evidence suggests that impaired cognitive function, specifically dysfunction in frontal cortical areas subserving executive functions, contributes to motor dysfunction in Parkinson’s disease. All prior studies, however, are correlational. We demonstrate that modulating DLPFC function with non-invasive brain stimulation causally impacts motor performance in people with PD. Administering an excitatory stimulation protocol (iTBS) improved motor performance relative to both an inhibitory stimulation protocol (cTBS) and stimulation of a control site, with the strongest effects emerging under dual-task conditions. Behavioral enhancement coincided with changes in functional connectivity between DLPFC and primary motor cortex. Inter-effector regions of motor cortex within the somato-cognitive action network (SCAN) showed widespread shifts in connectivity, including altered coupling with DLPFC and striatal areas implicated in cognition and action. These results provide causal evidence demonstrating that cognitive control circuitry influences motor performance in Parkinson’s disease.

Excitatory stimulation of DLPFC improved motor performance in a precision force-tracking task and effects emerged preferentially under dual-task conditions when cognitive demands were high. Improved performance cannot be explained by a shift in priority whereby participants focused more on the force-tracking task at the expense of cognitive performance, as we did not observe any decrement in performance on the concurrent N-back task following stimulation.^13^ These results are in line with prior suggestions that an Attentional-Motor Interface links prefrontal cortical regions with basal ganglia and subcortical cholinergic systems to support the integration of cognitive and motor processes in PD.^45^ Several studies suggested that intact Attentional-Motor Interface function enables compensatory recruitment of cognitive resources to maintain motor performance and that degeneration of these circuits, particularly within cholinergic pathways, contributes to gait dysfunction, freezing, and falls.^11,45,65,66^ By enhancing DLPFC activity, we provide converging causal evidence for the claim that increased engagement of prefrontal regions can improve motor performance in PD.

Improved motor performance following excitatory stimulation over DLPFC coincides with reductions in functional connectivity not only between the stimulation site and primary motor cortex, but also within the recently described SCAN. Ren et al.^16^ suggested that increased connectivity within SCAN is a pathophysiologic hallmark of parkinsonism. It is currently unclear whether the reduced connectivity we observed reflects greater efficiency in communication or reduced maladaptive coupling. Although increased functional connectivity is often interpreted as a marker of compensation, hyperconnectivity between cortico-cortico and cortico-subcortical networks is a reliable marker of dysfunction.^20,67–69^ Within this framework, the reductions we observed following iTBS are consistent with a relative normalization of network connectivity rather than a loss of functional integration, a result consistent with Ren et al.’s^16^ conclusion that heightened inter-SCAN connectivity is a marker of PD. Our findings support a model in which Parkinson’s disease reflects aberrant coupling between cognitive and motor areas rather than isolated dysfunction in motor networks. In this context, the observed decreases in DLPFC–SCAN and SCAN–striatal connectivity following iTBS suggest that targeting cognitive control regions normalize what may be pathological network interactions that, in turn, improve motor function.

We found limited evidence that inhibitory stimulation led to reliable effects relative to control stimulation. If cognitive networks play a compensatory role in supporting motor performance, their inhibition might worsen motor performance. Participants’ performance on the precision force tracking task used here was quite poor at baseline and may have been near floor levels of performance. There may have been less room for inhibitory stimulation to have a detrimental effect on motor performance. However, we did find some evidence that inhibitory stimulation led to poor performance on the concurrent n-back task. It is possible that following inhibitory stimulation, participants shifted attentional priority toward the motor task to maintain motor performance at the expense of cognitive performance. In either case, the inclusion of a condition with inhibitory stimulation provides a useful control condition to confirm that specifically excitatory stimulation of DLPFC activity leads to beneficial effects on motor performance.

Most studies examining the effects of non-invasive brain stimulation in Parkinson’s disease targeted cortical motor regions such as M1 or supplementary motor area.^70–74^ Behavioral outcomes are mixed, where similar stimulation sites and protocols lead to inconsistent results.^71,75^ Few studies have stimulated DLPFC to assess changes in motor behavior^76^ in Parkinson’s disease with mixed results and many null findings.^30,77,78^ We recently suggested that inconsistent behavioral outcomes following brain stimulation may arise from a reliance on simple motor tasks to evaluate performance and a narrow focus on primary motor areas as stimulation targets.^79^ For example, many studies^77,78,80^ use hand movements, tapping tasks, and pegboard tasks as an assay of motor behavior, but these metrics of performance may be too crude to detect subtle changes in behavior. Our findings highlight the value of more complex motor assessments and suggest that targeting non-motor regions may prove more effective in enhancing performance.

We cannot rule out that DLPFC stimulation affects a larger network of brain regions. We did not observe any significant effects of stimulation on univariate activity or functional connectivity in whole-brain analyses (data not shown). This may be due to our relatively modest sample size, which did not afford the statistical power necessary to overcome the stringent multiple comparisons correction necessary. It is possible that altering DLPFC activity leads to more widespread changes in cortical and subcortical networks previously implicated in PD. Future work with larger samples and whole-brain approaches will be important for confirming the generalizability of these findings and further characterizing how stimulation of cognitive control networks influences motor function in Parkinson’s disease.

By demonstrating that excitatory DLPFC stimulation can improve motor performance during cognitively demanding situations, our findings suggest that cognitive control circuitry could be a potential target for treating motor dysfunction in Parkinson’s disease. Our results were robust after a single session of stimulation, but it is unclear how long the beneficial effects of stimulation last, as we only assessed performance during the hour following stimulation. Future work is necessary to determine whether larger and more durable benefits can be achieved. We recently outlined how multi-day stimulation protocols and other modifications such as state-dependent neuromodulation and network-targeted stimulation might be advantageous in using TMS to improve cognitive and motor dysfunction in PD.^79^ Repeated stimulation sessions, including accelerated paradigms that deliver multiple sessions per day, demonstrated substantial and sustained clinical benefits in other disorders. This prior work may be a model for how TMS might be used for therapeutic benefit in PD as it is possible that repeated bouts of excitatory stimulation of DLPFC will induce long-lasting benefits.

Our findings demonstrate that excitatory stimulation of DLPFC can improve motor performance in Parkinson’s disease and provide causal evidence that cognitive control networks contribute to voluntary movement control. These behavioral improvements were accompanied by changes in functional connectivity across distributed cognitive-motor networks, supporting the view that motor impairments in Parkinson’s disease arise not solely from dysfunction within traditional motor circuits, but also from disruptions in a distributed cognitive-motor network spanning both cortical and subcortical regions.

## Data availability

Behavioral datasets underlying the statistical analyses reported in this study will be publicly available through the Open Science Framework (OSF) at https://osf.io/ahctp/. Owing to the large size of the raw MRI datasets and associated storage constraints, raw neuroimaging data are not included in the public repository but may be made available by the corresponding author upon reasonable request.

## Acknowledgements

We would like to thank the Functional MRI Laboratory at the University of Michigan. We would like to thank Jacob Sellers and Quynh Nguyen for helpful comments and discussion. We thank Ritika Tiwary and Julian Singleton for help with data collection.

